# Selection shapes the genomic landscape of introgressed ancestry in a pair of sympatric sea urchin species

**DOI:** 10.1101/2023.12.01.566927

**Authors:** Matthew R. Glasenapp, Grant H. Pogson

## Abstract

A growing number of recent studies have demonstrated that introgression is common across the tree of life. However, we still have a limited understanding of the fate and fitness consequence of introgressed variation at the whole-genome scale across diverse taxonomic groups. Here, we implemented a phylogenetic hidden Markov model to identify and characterize introgressed genomic regions in a pair of well-diverged, non-sister sea urchin species: *Strongylocentrotus pallidus* and *S. droebachiensis*. Despite the old age of introgression, a sizable fraction of the genome (1% - 5%) exhibited introgressed ancestry, including numerous genes showing signals of historical positive selection that may represent cases of adaptive introgression. One striking result was the overrepresentation of hyalin genes in the identified introgressed regions despite observing considerable overall evidence of selection against introgression. There was a negative correlation between introgression and chromosome gene density, and two chromosomes were observed with considerably reduced introgression. Relative to the non-introgressed genome-wide background, introgressed regions had significantly reduced nucleotide divergence (*d*_XY_) and overlapped fewer protein-coding genes, coding bases, and genes with a history of positive selection. Additionally, genes residing within introgressed regions showed slower rates of evolution (*d*_N_, *d*_S_, *d*_N_/*d*_S_) than random samples of genes without introgressed ancestry. Overall, our findings are consistent with widespread selection against introgressed ancestry across the genome and suggest that slowly evolving, low-divergence genomic regions are more likely to move between species and avoid negative selection following hybridization and introgression.

## Introduction

Advances in genome sequencing have revealed that many species hybridize with close relatives and share alleles through introgression. However, relatively little is known about the fate and fitness consequence of introgressed variation at the whole-genome scale. Although it is well established that introgression often facilitates adaptation (Song et al. 2011; The Heliconius Genome Consortium 2012; Huerta-Sánchez et al. 2014; Lamichhaney et al. 2015; Arnold et al. 2016), studies documenting adaptive introgression often focus on phenotypes that were known *a priori* to have been involved in adaptation (Martin and Jiggins 2017). Despite enthusiasm about the potential for introgression to introduce new alleles already tested by selection at a frequency higher than mutation (Hedrick 2013; Martin and Jiggins 2017), there remain few estimates of the proportion of introgressed ancestry fixed by positive selection across entire genomes.

A promising approach to understanding the overall fitness consequence of introgression involves identifying and characterizing genomic regions showing introgressed ancestry. Genome-wide scans for introgression have demonstrated that the distribution of introgressed haplotypes across the genome, termed the genomic landscape of introgression, is highly heterogeneous due to the combined effects of natural selection and recombination. Most introgressed variation is thought to be deleterious and removed by selection because it either (i) is maladapted to the recipient’s ecology (i.e., ecological selection; McBride and Singer 2010; Arnegard et al. 2014; Cooper et al. 2018), (ii) causes negative epistasis in the genomic background of the recipient (i.e., hybrid incompatibilities; Orr 1995), or (iii) imposes a genetic load on the recipient if the donor has a smaller effective population size (i.e., hybridization load; Harris and Nielsen 2016; Juric et al. 2016; but see Kim et al. 2018). Selection against introgression should lead to a positive correlation between introgression and recombination because recombination weakens the strength of selection acting on introgressed variation by breaking up long introgression tracts with multiple linked, deleterious alleles and distributing introgressed ancestry more evenly among individuals (Barton 1983; Harris and Nielsen 2016; Veller et al. 2023). Numerous empirical studies have demonstrated a positive correlation between introgression and the local recombination rate (Brandvain et al. 2014; Schumer et al. 2016; Ravinet et al. 2018; Martin et al. 2019; Calfee et al. 2021; Ravinet et al. 2021). However, few organismal groups have been studied, and some notable exceptions exist, including a negative correlation between local recombination rate and admixed ancestry in *Drosophila melanogaster* (Pool 2015; Corbett-Detig and Nielsen 2017).

Gene density is also thought to influence the distribution of introgressed ancestry, as the strength of selection depends on the density of selected sites (Barton 1983; Barton and Bengtsson 1986; Martin and Jiggins 2017). Reduced rates of introgression near functionally important elements have been found in diverse groups, such as humans (Sankararaman et al. 2014; Juric et al. 2016; Sankararaman et al. 2016; Petr et al. 2019), house mice (Teeter et al. 2008; Janoušek et al. 2015), *Histoplasma* fungi (Maxwell et al. 2018), wild strawberries (Feng et al. 2023), and *Xiphophorus* swordtails (Schumer et al. 2016). It is important to note that gene density and recombination rate are often not independent, further complicating the interpretation of introgression patterns (Martin and Jiggins 2017). For example, a positive correlation between recombination rate and gene density may lead to higher retention of introgression in gene-dense regions than expected (Schumer et al. 2016; Baker et al. 2017; Schumer et al. 2018; Moran et al. 2021).

Within the protein-coding portion of the genome, introgressed variation should be less common at genes with high divergence or faster rates of adaptive evolution because they are more likely to underlie locally adapted phenotypes or hybrid incompatibilities. However, few empirical studies have tested this prediction. Among modern humans, introgressed Neanderthal ancestry is negatively correlated with fixed differences between humans and Neanderthals, consistent with introgressed variation having negative fitness consequences in high-divergence regions (Vernot and Akey 2014). Conversely, Schumer et al. (2016) found that introgressed loci in *Xiphophorus* swordtails had higher divergence than non-introgressed loci due to reduced selective constraint. Characterizing patterns of introgression across a broader range of taxonomic groups is needed to better understand the factors influencing the distribution of introgression along genomes.

The strongylocentrotid family of sea urchins is a compelling group for characterizing the genomic landscape of introgression. Extensive introgression has occurred among the strongylocentrotid urchins (Glasenapp and Pogson 2023), and the purple sea urchin, *Strongylocentrotus purpuratus* (Stimpson), has a well-annotated reference genome and a long history of use as a model organism in fertilization and development studies. The selective histories of the single-copy protein-coding genes in this family have been formally characterized (Kober and Pogson 2017), providing valuable context for interpreting introgression patterns. The massive effective population sizes of sea urchins should lead to considerably more efficient selection on introgressed variation than most model systems studied thus far. Additionally, their high amounts of recombination (Brennan et al. 2019) and lack of population structure (Palumbi and Wilson 1990; Palumbi and Kessing 1991) should promote the retention of introgressed ancestry. Several studies have documented introgression between a pair of recently-diverged, non-sister taxa that co-occur and hybridize in both the North Pacific and North Atlantic: *S. pallidus* and *S. droebachiensis* (Addison and Hart 2005; Harper and Hart 2007; Addison and Pogson 2009; Pujolar and Pogson 2011; Glasenapp and Pogson 2023). However, the genomic regions showing signals of introgression have yet to be characterized.

To better understand the fitness consequence of introgression in strongylocentrotid sea urchins, we characterized genomic regions exhibiting introgressed ancestry between *S. pallidus* and *S. droebachiensis* and asked whether these regions show any nonrandom patterns compared to the genome-wide background. We predicted that introgressed regions would have lower gene density, divergence, and rates of evolution than the genome-wide background and that genes with a history of positive selection would show reduced rates of introgression. We further looked for potential cases of adaptive introgression and tested whether introgressed genes were enriched for any gene families or functional categories.

## Results

To identify genomic regions supporting introgression between non-sister taxa *S. pallidus* and *S. droebachiensis*, we applied the phylogenetic hidden Markov model PhyloNet-HMM (Liu et al. 2014) to pseudo-haploid multi-species multiple sequence alignments of the 21 largest scaffolds in the *S. purpuratus* reference genome assembly (Figure 1). The multiple sequence alignments were constructed from hard-filtered genotypes of each species in the rooted triplet (*Hemicentrotus pulcherrimus*, (*Strongylocentrotus pallidus*, (*S. droebachiensis*, *S. fragilis*))). The genotypes were obtained by mapping paired-end sequencing reads of each species to the *S. purpuratus* reference genome with bwa-mem2 (Vasimuddin et al. 2019) and calling and genotyping variants following GATK’s Best Practices (Van der Auwera et al. 2013). The 21 largest scaffolds in the Spur_5.0 assembly correspond to the 21 *S. purpuratus* chromosomes (2n=42) and represent 90% of the bases in the 922 Mb assembly. The PhyloNet-HMM model walks along each chromosome, identifies changes in the underlying genealogy, and outputs posterior probabilities of having evolved by each parent tree (i.e., species tree, introgression tree) for each site in the multiple sequence alignment (Liu et al. 2014). The model accounts for both convergence and incomplete lineage sorting (ILS) by employing a finite-sites model and allowing for changes in gene trees within each parent tree (Liu et al. 2014; Liu et al. 2015; Schumer et al. 2016). We ran PhyloNet-HMM 100 times on each scaffold and averaged the posterior probabilities across runs to avoid the effects of reaching local optima during hill climbing (per suggestion by PhyloNet-HMM developer Qiqige Wuyun). To infer introgression tracts, we applied a posterior probability threshold for introgression of 90% and recorded the genomic coordinates of consecutive sites with posterior probabilities at or above this threshold. We also proceeded with a less stringent dataset at the 80% posterior probability threshold for comparison, given the conservative nature of our test (see Discussion) and the small size of the dataset identified at the 90% threshold. In both datasets, the inferred introgression tracts were filtered if they had a mean coverage depth less than 5x or greater than 100x and trimmed if they overlapped a gap of more than 25kb between adjacent genotypes (including invariant sites).

**Figure 1.**
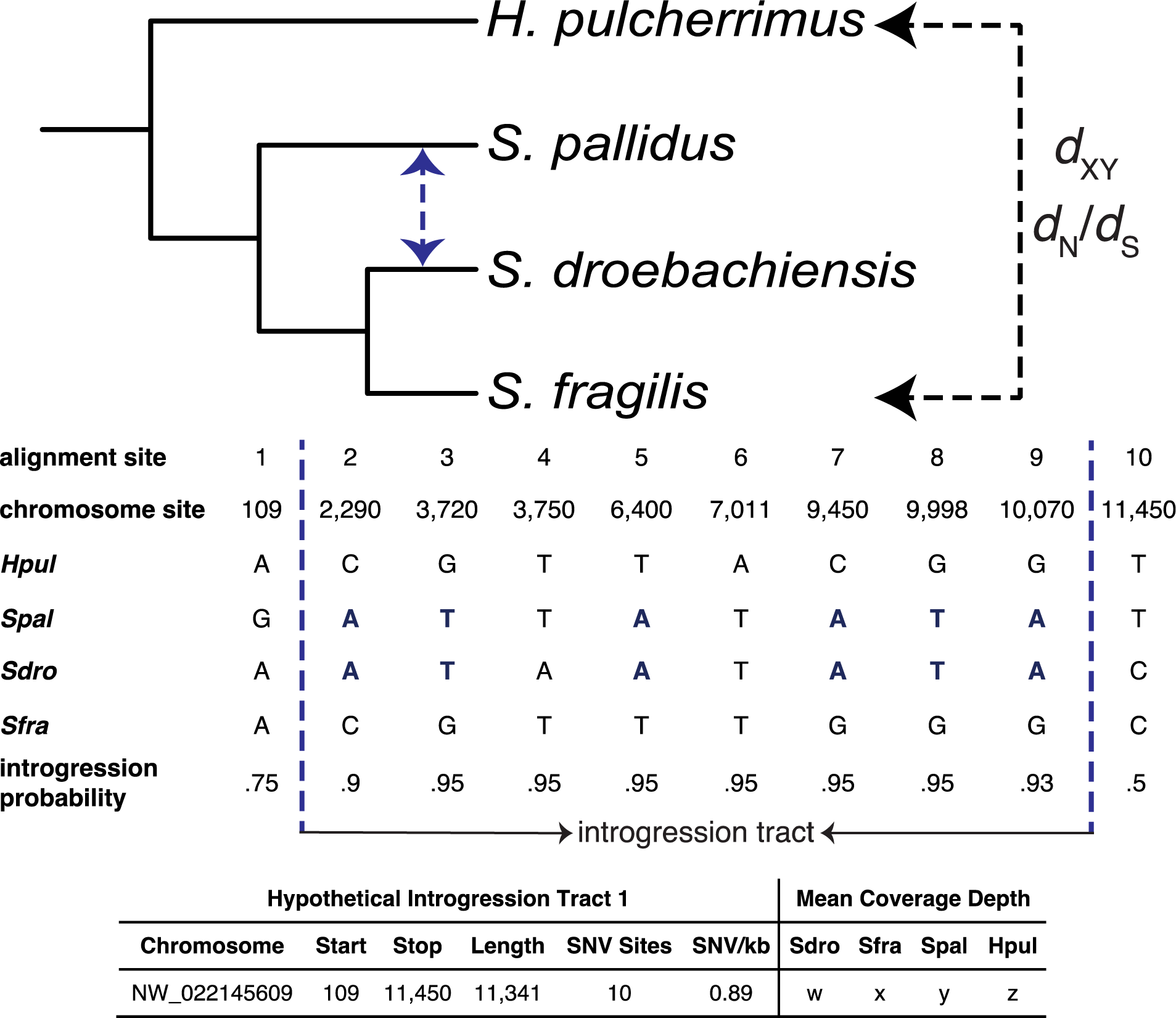
Schematic of the study design. Introgressed genomic regions between *S. pallidus* and *S. droebachiensis* were identified using PhyloNet-HMM. PhyloNet-HMM outputs posterior probabilities of introgression for each site in a multiple sequence alignment. Introgression tracts were inferred by recording the genomic coordinates of consecutive sites with posterior probabilities at or above two different thresholds (90%, 80%). For each introgression tract, coverage depth metrics were estimated, and a gene tree was reconstructed for the region. For introgression tracts longer than 10kb, estimates of *H. pulcherrimus* - *S. fragilis d*_XY_ were made and compared to estimates obtained from genomic regions with high support for the species tree. All genes overlapping introgression tracts were identified. Estimates of *d*_N_/*d*_S_ from sequence alignments of *H. pulcherrimus* and *S. fragilis* were made for genes with more than half of their bases declared introgressed and compared to estimates for non-introgressed genes. *S. pallidus* and *S. droebachiensis* were not used in the *d*_XY_ or *d*_N_/*d*_S_ estimates because introgression in either direction could confound the estimates.

At the 90% posterior probability threshold, we identified 4,855 introgression tracts (≥2 bases), with 164 exceeding 10kb. Tracts greater than 10kb in length had mean and median lengths of 22,850 and 16,595 base pairs, respectively (Supplementary Figure S1). The coverage depth and breadth metrics for introgression tracts were similar to the genome-wide averages (Table 1, Supplementary Table S2). The percent of bases introgressed, both overall and in coding regions, was 1%.

**Table 1.** Summary of DNA sequencing and coverage.

| Species | % Bases Covered by 5 reads |  |  | Mean Coverage Depth |  |  |
| --- | --- | --- | --- | --- | --- | --- |
|  | Whole Genome<sup>a</sup> | Coding<sup>b</sup> | Introgression Tracts<sup>c</sup> | Whole Genome<sup>d</sup> | Coding<sup>e</sup> | Introgression Tracts<sup>f</sup> |
| *Sdro* | 62 | 90 | 66 | 24.7x | 52.9x | 25.4x |
| *Sfra* | 69 | 91 | 73 | 32.1x | 54.8x | 32.7x |
| *Spal* | 57 | 85 | 61 | 11.9x | 17.3x | 13.5x |
| *Hpul* | 53 | 83 | 54 | 24.5x | 54.2x | 23.5x |

<sup>a</sup>Percentage of bases in the *S. purpuratus* reference genome covered with at least five reads.
<sup>b</sup>Percentage of coding bases in the *S. purpuratus* reference genome covered with at least five reads.
<sup>c</sup>Percentage of bases in the introgression tracts identified at the 90% posterior probability threshold covered with at least five reads.
<sup>d</sup>Mean genome-wide coverage depth of the *S. purpuratus* reference genome
<sup>e</sup>Mean coverage depth for coding sequences in the *S. purpuratus* genome assembly
<sup>f</sup>Mean coverage depth for the introgression tracts identified at the 90% posterior probability threshold.
Species abbreviations: *Sdro* - *S. droebachiensis*; *Sfra* - *S. fragilis*; *Spal* - *S. pallidus*; *Hpul* - *H. pulcherrimus*

When the posterior probability threshold was lowered to 80%, the total number of tracts increased to 17,037, with 953 exceeding 10kb. The mean and median tract lengths for 10kb tracts (22,897, 17,236 base pairs) remained similar to those at the 90% threshold. The percentage of bases introgressed rose to 5% overall and 6% in coding regions. These estimates of the proportion of the genome introgressed (1 - 5 %) align with previous estimates for *S. pallidus* and *S. droebachiensis* (Glasenapp and Pogson, 2023). Summary statistics for the introgression tracts at both probability thresholds are provided in Supplementary Table S3, and information on all introgression tracts can be found in Supplementary Tables S4 and S5. Additionally, Figure 2 and Supplementary Figure S2 depict the locations of the 10 kb introgression tracts along scaffolds.

**Figure 2.**
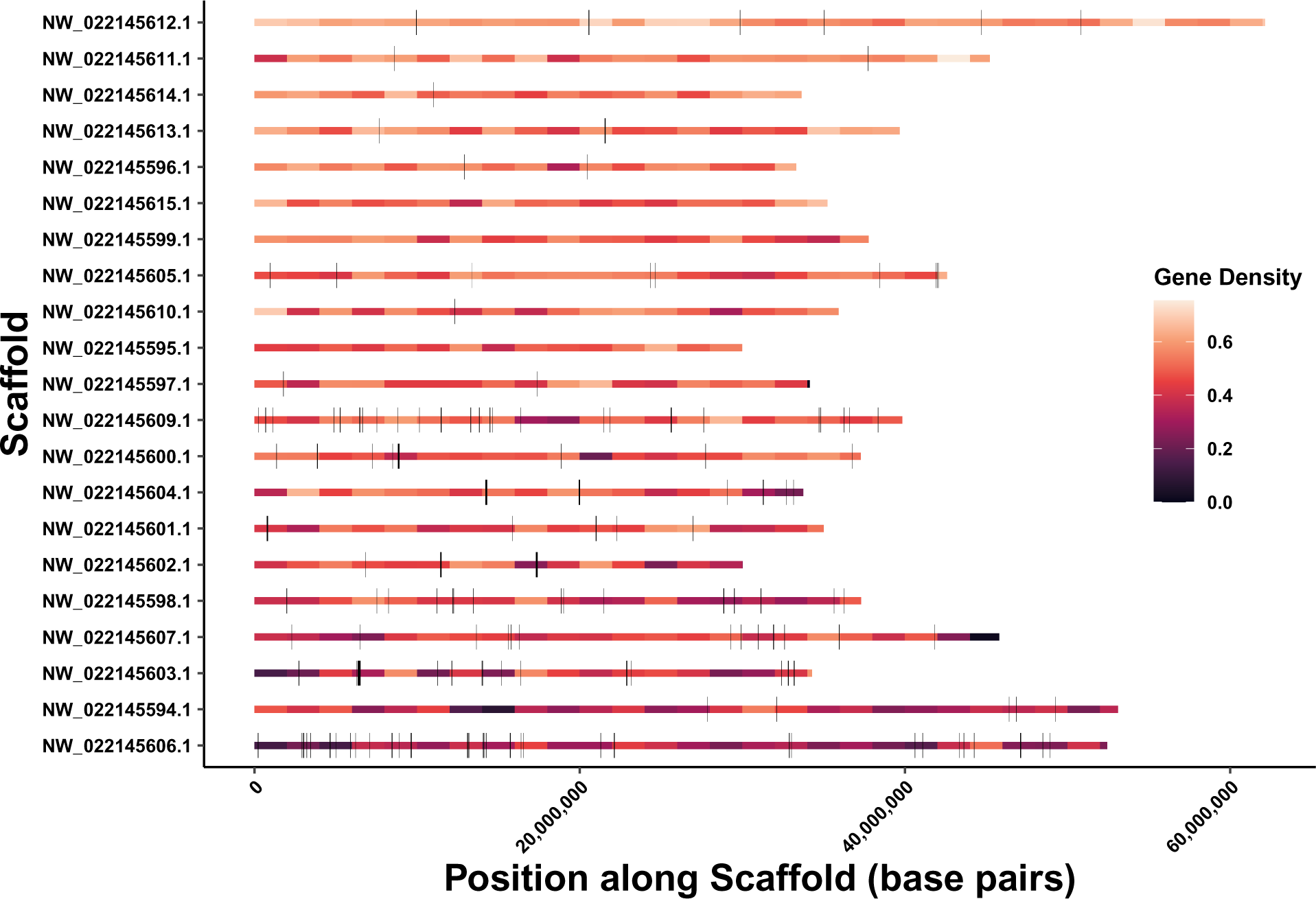
The 164 introgression tracts greater than 10 kb in length by chromosome (posterior probability > 90%). The introgression tracts are displayed as black rectangles along the chromosomes. The chromosomes are ordered by gene density (descending). The chromosomes are colored by gene density in windows of 2Mb.

There was a significantly negative association between the percent scaffold introgressed and scaffold-wide gene density (Supplementary Figure S3). The scaffold with the highest percent introgressed and most 10kb introgression tracts (NW_022145606.1) also had the lowest gene density (Supplementary Table S6). Unexpectedly, two scaffolds (NW_022145615.1, NW_022145595.1) did not have any sites that crossed the 90% posterior probability threshold for introgression (Supplementary Table S6). There were no discernible features of these two chromosomes that would lead to reduced power to detect introgression. Both had high site density (Supplementary Table S6) and although NW_022145595.1 is the shortest scaffold at 30 Mb, fifteen of the 21 scaffolds were between 30 and 40 Mb in length. To determine the probability of having two chromosomes without 10 kb introgression tracts due to chance, we divided the genome into non-overlapping 10kb blocks and selected 164 blocks at random 10,000 times, recording the frequency of one or more chromosomes not being represented. The number of times a single chromosome was not represented was 154 in 10,000 (0.015). The number of times two chromosomes were not represented was 1 in 10,000 (0.0001). At the 80% posterior probability threshold, all chromosomes had 10kb introgression tracts, ranging from 9-127 (Supplementary Table S7). Consistent with the results at the 90% threshold level, scaffolds NW_022145615.1 and NW_022145595.1 had the fewest percent of introgressed sites (1.4%, 1.5%) and 10kb introgression tracts (9, 10), while scaffold NW_022145606.1 again had the highest proportion of introgressed sites (11.5%) and the most 10kb introgression tracts (127) (Supplementary Table S7).

To characterize the properties of introgressed genomic regions, we compared estimates of absolute nucleotide divergence (*d*_XY_), gene density, coding base density, the rate of nonsynonymous substitutions (*d*_N_), the rate of synonymous substitutions (*d*_S_), and the nonsynonymous to synonymous substitution rate ratio (*d*_N_/*d*_S_) for introgressed regions and genes to the non-introgressed genome-wide background. To avoid the confounding effect of introgression between *S. pallidus* and *S. droebachiensis* on *d*_XY_ estimates, we used *H. pulcherrimus* and *S. fragilis*, who have experienced little-to-no introgression. We implemented a bootstrap comparison of means to compare *d*_XY_ between introgression tracts and the non-introgressed genome-wide background. First, we calculated *H. pulcherrimus* - *S. fragilis d*_XY_ for all introgression tracts, and a random sample of regions of the same number and size confidently called for the species tree. We then pooled all *d*_XY_ values, bootstrap resampled the pool in pairs 100,000 times, and calculated the difference in mean *d*_XY_ between bootstrapped pairs to generate the distribution of differences in means expected if there were no difference between mean introgressed *d*_XY_ and mean non-introgressed *d*_XY_. We then compared the true difference in mean *d*_XY_ between the introgressed and non-introgressed regions to the null distribution to calculate a p-value. We found that introgressed regions had lower divergence (*d*_XY_) than the genome-wide background at both posterior probability thresholds (p<0.0001; Table 2, Figure 4, Supplementary Figure S4).

**Table 2.** Genomic features of the distribution of 10 kb introgression tracts and the genome-wide background at the 90% posterior probability threshold. Means are shown for the introgression tracts. The 95% confidence intervals are shown for the genome-wide background.

| | $d_{XY}$ | $d_N$ | $d_S$ | $d_N/d_S$ | Genes / Mb | Percent Coding | PSGs<sup>a</sup>/ Mb |
| --- | --- | --- | --- | --- | --- | --- | --- |
| **Introgression Tracts/Genes** | 0.041 | 0.001 | 0.05 | 0.19 | 17.6 | 3.75 | 0.68 |
| **Genome-Wide Background** | 0.048 - 0.054 | 0.011 - 0.019 | 0.042 - 0.069 | 0.25 - 0.45 | 28.6 - 28.9 | 5.19 - 5.23 | 1.58 - 1.65 |

<sup>a</sup>Positively Selected Genes: Genes with significant sites tests of positive selection across the stronglycentrotid family.

To compare the rate of evolution (*d*_N_, *d*_S_, and *d*_N_/*d*_S_) between introgressed and non-introgressed regions, the same procedure used in the *d*_XY_ analysis was repeated for genes with more than half of their bases declared introgressed and a random set of the same number of genes with more than half of their bases confidently called for the species tree. Genes were filtered if they had a mean coverage depth less than 10x or greater than 100x, had fewer than 75% of their coding bases covered by one read or fewer than 50% by ten reads, or contained stop codons. We estimated *d*_N_, *d*_S_, and *d*_N_/*d*_S_ using codeml M0 of PAML (Yang 2007). At the 90% posterior probability, introgressed genes had lower *d*_N_ (p=0.03), *d*_S_ (p=0.34), and *d*_N_/*d*_S_ (p<0.0001) than non-introgressed genes (Table 2, Figure 4, Supplementary Table S8). The relationships remained the same at the 80% posterior probability threshold, with the difference in mean *d*_S_ becoming significant (p=0.001; Supplementary Table S8, Supplementary Figure S5).

We then compared the number of overlapping protein-coding genes, number of overlapping coding bases, and number of overlapping genes with a history of positive selection between the introgression tracts and the non-introgressed genome-wide background. Following the approach of Schumer (2016), we generated distributions of the counts of overlapping genes, coding bases, and positively selected genes for introgression tracts by bootstrap resampling the 10kb introgression tracts with replacement 1,000 times and counting the number of overlapping features. We calculated the mean and standard deviation for each metric. We then compared these means to null distributions created by randomly permuting intervals of the same number and size as the introgression tracts into the genomic regions confidently called for the species tree 1,000 times and counting the number of overlapping genes, coding bases, and positively selected genes. Introgression tracts overlapped fewer protein-coding genes, coding bases, and genes with a history of positive selection at both posterior probability thresholds (Table 2, Figure 4, Supplementary Table S8, Supplementary Figure S5).

We further identified all genes overlapping introgression tracts at both posterior probability thresholds (Supplementary Table S9, S10) using gene models from the latest *S. purpuratus* genome assembly (Spur_5.0). At the 90% posterior probability threshold, 50 protein-coding genes had all their bases declared introgressed, and another 102 had more than half of their bases introgressed. A total of 2,055 genes overlapped an introgression tract by at least two bases. One noteworthy pattern was that many different hyalin genes had bases declared introgressed. Hyalin, an extracellular matrix glycoprotein, is the major component of the hyaline layer, an extraembryonic matrix serving as a cell adhesion substrate during development (McClay and Fink 1982; Wessel et al. 1998). At the 90% threshold, five unique hyalin genes had bases introgressed (LOC578156, LOC752152, LOC373362, LOC100891695, LOC100891850), and an additional four hyalin genes showed introgression at the 80% threshold (LOC578713, LOC586885, LOC576524, LOC578967). Coverage depth for all but one of these genes (LOC578967) was in the range expected if they were single-copy across the four species analyzed (Supplementary Table S11). The high number of hyalin genes observed may not be unexpected, given that there are 21 hyalin genes in the *S. purpuratus* assembly. To test whether there were more occurrences of hyalin than expected by chance, we randomly sampled the same number of genes as the number of introgressed genes from the set of all genes on the 21 chromosomes 1,000 times and recorded the number of hyalin occurrences. There were 3.3x more hyalin genes in the introgressed set than expected due to chance at the 90% threshold (95% confidence interval: 1.4 - 1.6) and 1.8x at the 80% threshold (95% confidence interval: 4.9 - 5.2). More occurrences of hyalin gene introgression may be expected than due to random chance if they are clustered near each other on chromosomes and/or do not segregate independently. However, the introgressed hyalin genes have nonoverlapping coordinates and occur across six different chromosomes, with the shortest gap between genes sharing a chromosome being > 400kb. Furthermore, in no case did a single introgression tract overlap more than one hyalin gene.

To identify other potential examples of adaptive introgression, we looked for overlap between genes with introgressed coding bases and the 1,008 strongylocentrotid single-copy orthologs with a history of positive selection previously identified by Kober and Pogson (2017). The positively selected genes (PSGs) had been previously identified by comparing the codon sites models M7 (Beta) *vs*. M8 (Beta plus ω) (Yang et al. 2000) using the CODEML program of the PAML Package (Yang, 2007). Branch-sites tests of positive selection were also used to identify lineage-specific episodes of adaptive evolution (Kober and Pogson 2017). At the 90% confidence level, three genes with significant sites tests across the family had more than half of their coding bases declared introgressed: arachidonate 5-lipoxygenase (Figure 3), helicase domino, and kinesin-II 95 kDa subunit (Table 3, Supplementary Table S12). There were 32 PSGs (3.2%) with at least 10% of their coding bases introgressed. At the 80% posterior probability threshold, there was a total of 24 PSGs (2.4%) with more than half of their coding bases declared introgressed (Supplementary Table S13), including six genes that had all their coding bases declared introgressed: arachidonate 5-lipoxygenase, 5-hydroxytryptamine receptor 6, transcription termination factor 1, MAK16 homolog, glutathione peroxidase-like, and 2’,3’-cyclic-nucleotide 3’-phosphodiesterase-like (Table 3). The maximum likelihood gene trees for all introgressed PSGs shown in Table 3 grouped *S. pallidus* and *S. droebachiensis* as sister taxa, except for glutathione peroxidase-like, which did not have enough high-quality variant sites for gene tree reconstruction. All candidate introgressed PSGs had high coverage depth in the range expected for single-copy orthologs, indicating that genotyping error likely did not contribute to the introgression signal (Table 3).

**Figure 3.**
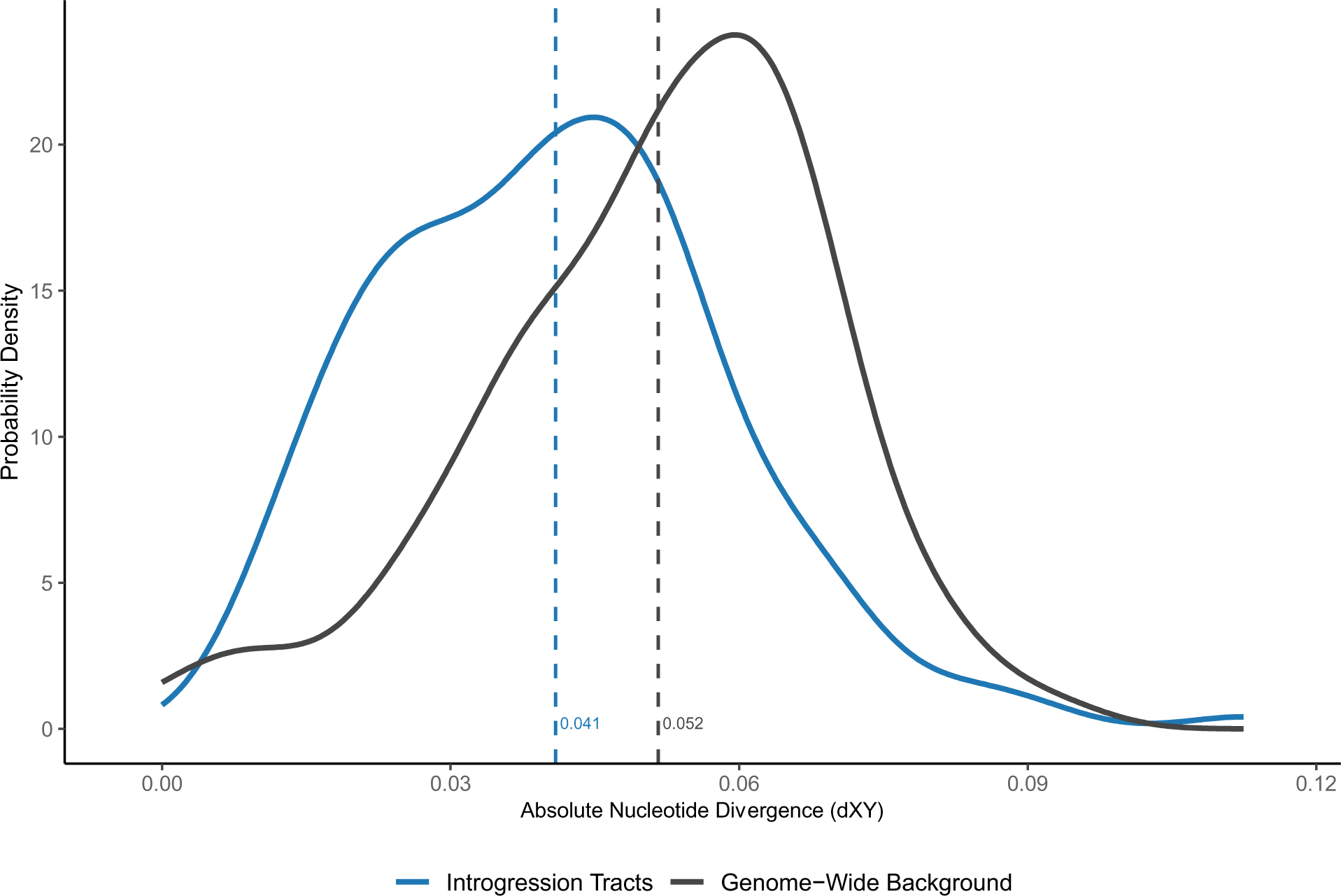
Absolute nucleotide divergence (*d*XY) between *H. pulcherrimus* and *S. fragilis* for the introgressed intervals vs. a random sample of non-introgressed intervals of the same number and length confidently called for the species tree by PhyloNet-HMM.

**Table 3.**
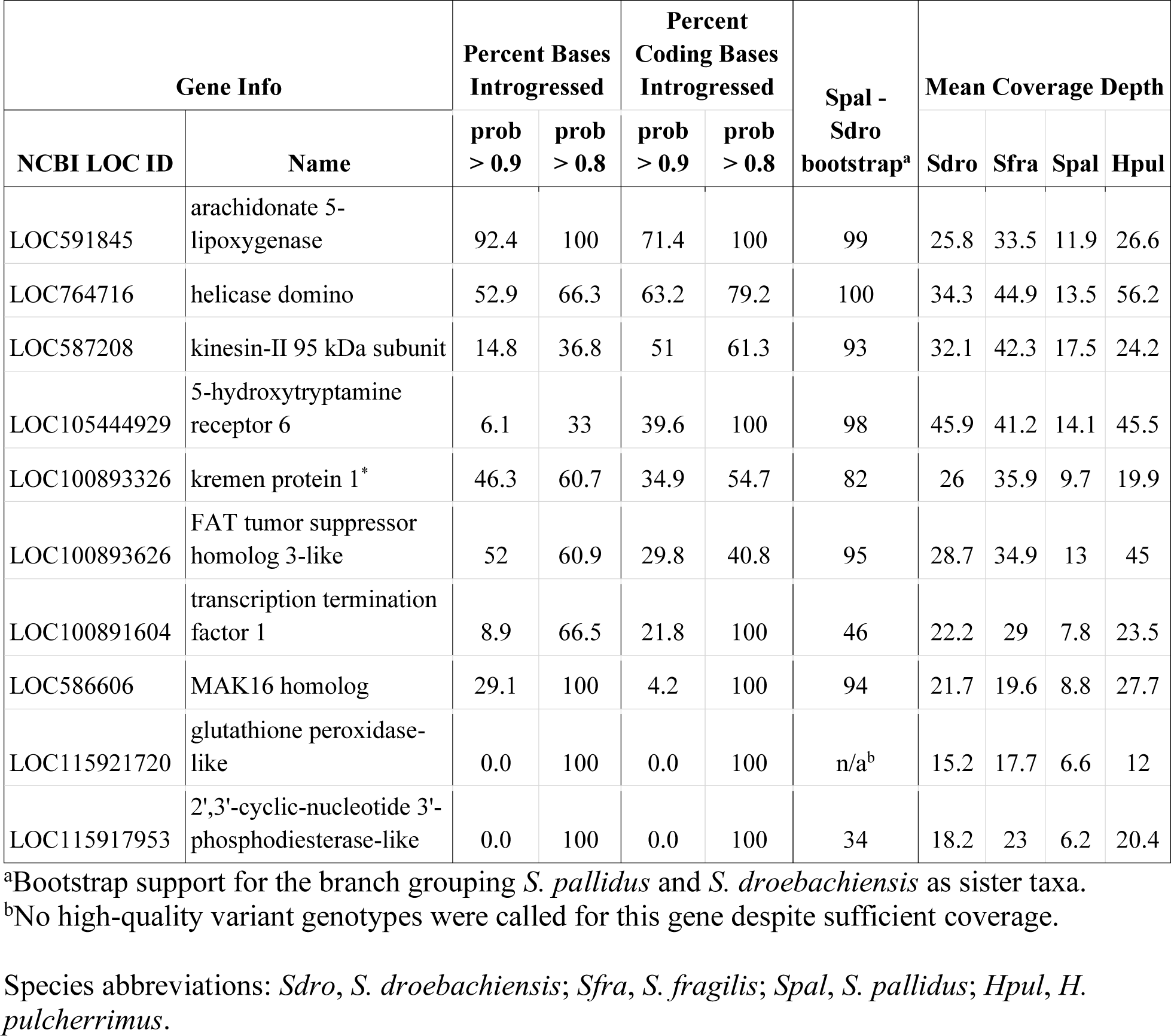
A selection of genes with a history of positive selection within the strongylocentrotid sea urchin family that overlapped introgression tracts. The list is organized by the percentage of coding bases introgressed at the 90% posterior probability threshold.

| Gene Info |  | Percent Bases Introgressed |  | Percent Coding Bases Introgressed |  | Spal - Sdro bootstrap<sup>a</sup> | Mean Coverage Depth |  |  |  |
| --- | --- | --- | --- | --- | --- | --- | --- | --- | --- | --- |
| NCBI LOC ID | Name | prob > 0.9 | prob > 0.8 | prob > 0.9 | prob > 0.8 |  | Sdro | Sfra | Spal | Hpul |
| LOC591845 | arachidonate 5-lipoxygenase | 92.4 | 100 | 71.4 | 100 | 99 | 25.8 | 33.5 | 11.9 | 26.6 |
| LOC764716 | helicase domino | 52.9 | 66.3 | 63.2 | 79.2 | 100 | 34.3 | 44.9 | 13.5 | 56.2 |
| LOC587208 | kinesin-II 95 kDa subunit | 14.8 | 36.8 | 51 | 61.3 | 93 | 32.1 | 42.3 | 17.5 | 24.2 |
| LOC105444929 | 5-hydroxytryptamine receptor 6 | 6.1 | 33 | 39.6 | 100 | 98 | 45.9 | 41.2 | 14.1 | 45.5 |
| LOC100893326 | kremen protein 1* | 46.3 | 60.7 | 34.9 | 54.7 | 82 | 26 | 35.9 | 9.7 | 19.9 |
| LOC100893626 | FAT tumor suppressor homolog 3-like | 52 | 60.9 | 29.8 | 40.8 | 95 | 28.7 | 34.9 | 13 | 45 |
| LOC100891604 | transcription termination factor 1 | 8.9 | 66.5 | 21.8 | 100 | 46 | 22.2 | 29 | 7.8 | 23.5 |
| LOC586606 | MAK16 homolog | 29.1 | 100 | 4.2 | 100 | 94 | 21.7 | 19.6 | 8.8 | 27.7 |
| LOC115921720 | glutathione peroxidase-like | 0.0 | 100 | 0.0 | 100 | n/a<sup>b</sup> | 15.2 | 17.7 | 6.6 | 12 |
| LOC115917953 | 2',3'-cyclic-nucleotide 3'-phosphodiesterase-like | 0.0 | 100 | 0.0 | 100 | 34 | 18.2 | 23 | 6.2 | 20.4 |

<sup>a</sup>Bootstrap support for the branch grouping *S. pallidus* and *S. droebachiensis* as sister taxa.
<sup>b</sup>No high-quality variant genotypes were called for this gene despite sufficient coverage.
Species abbreviations: *Sdro*, *S. droebachiensis*; *Sfra*, *S. fragilis*; *Spal*, *S. pallidus*; *Hpul*, *H. pulcherrimus*.

We also looked for overlap between introgressed genes and genes with significant branch-sites tests on the *S. pallidus* and *S. droebachiensis* terminal branches, indicating lineage-specific episodes of adaptive protein evolution (Yang 2005; Zhang et al. 2005). Four genes with significant branch-sites tests on the *S. pallidus* terminal branch had an appreciable proportion of their coding bases introgressed at the 90% posterior probability threshold: kremen protein 1, arylsulfatase, sodium- and chloride-dependent neutral and basic amino acid transporter B(0+), and fibrosurfin-like (Table 4). Two genes with significant branch-sites test on the *S. droebachiensis* terminal branch had coding bases introgressed at the 90% threshold, PHD finger protein 8, and structural maintenance of chromosomes 1A (Table 4). All introgressed genes with significant branch-sites tests in Table 4 supported an *S. pallidus* and *S. droebachiensis* sister relationship with an average bootstrap support of 85%. Additional genes with significant branch-sites tests and introgressed bases are available in Supplementary Tables S14-S17.

**Figure 4.**
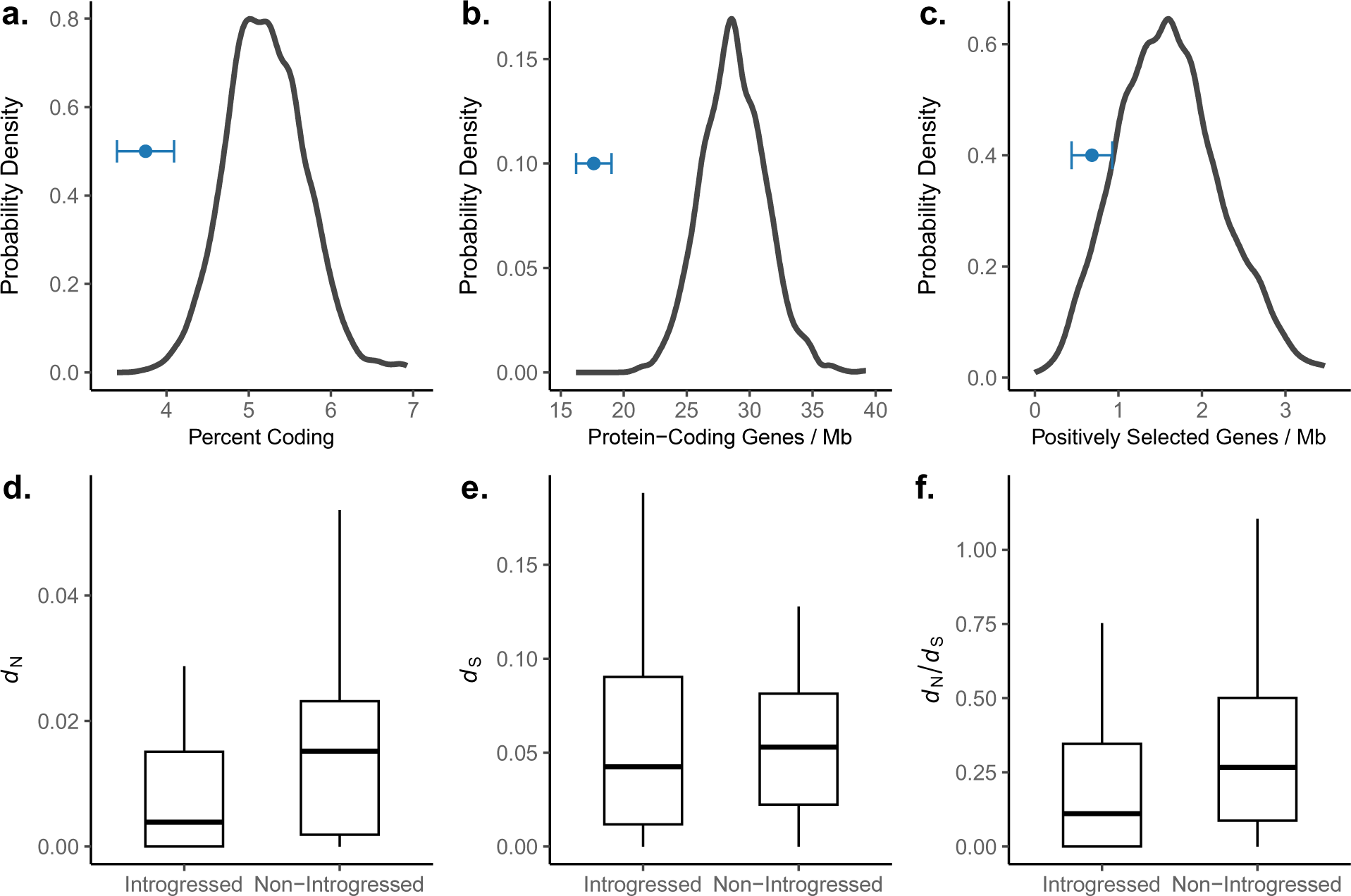
Properties of introgressed regions and genes relative to random non-introgressed genes representative of the genome-wide background. (a) The percentage of bases that are coding for the introgression tracts (blue) is lower than the genome-wide background. (b) The number of overlapping protein coding genes, standardized by the combined number of introgressed bases in Mb, is lower than the genome-wide background. (c) The number of overlapping positively selected genes, standardized by the number of bases in the interval files, is lower for introgression tracts than the genome-wide background. Errors bars in (a-c) represent the standard deviation. (d-f) *d*_N_, *d*_S_, and *d_N_*/*d*_S_ are lower for introgressed genes than non-introgressed genes. *d*_N_, *d*_S_, and *d_N_*/*d*_S_ were estimated on protein-coding alignments of *H. pulcherrimus* and *S. fragilis*.

**Figure 5.**
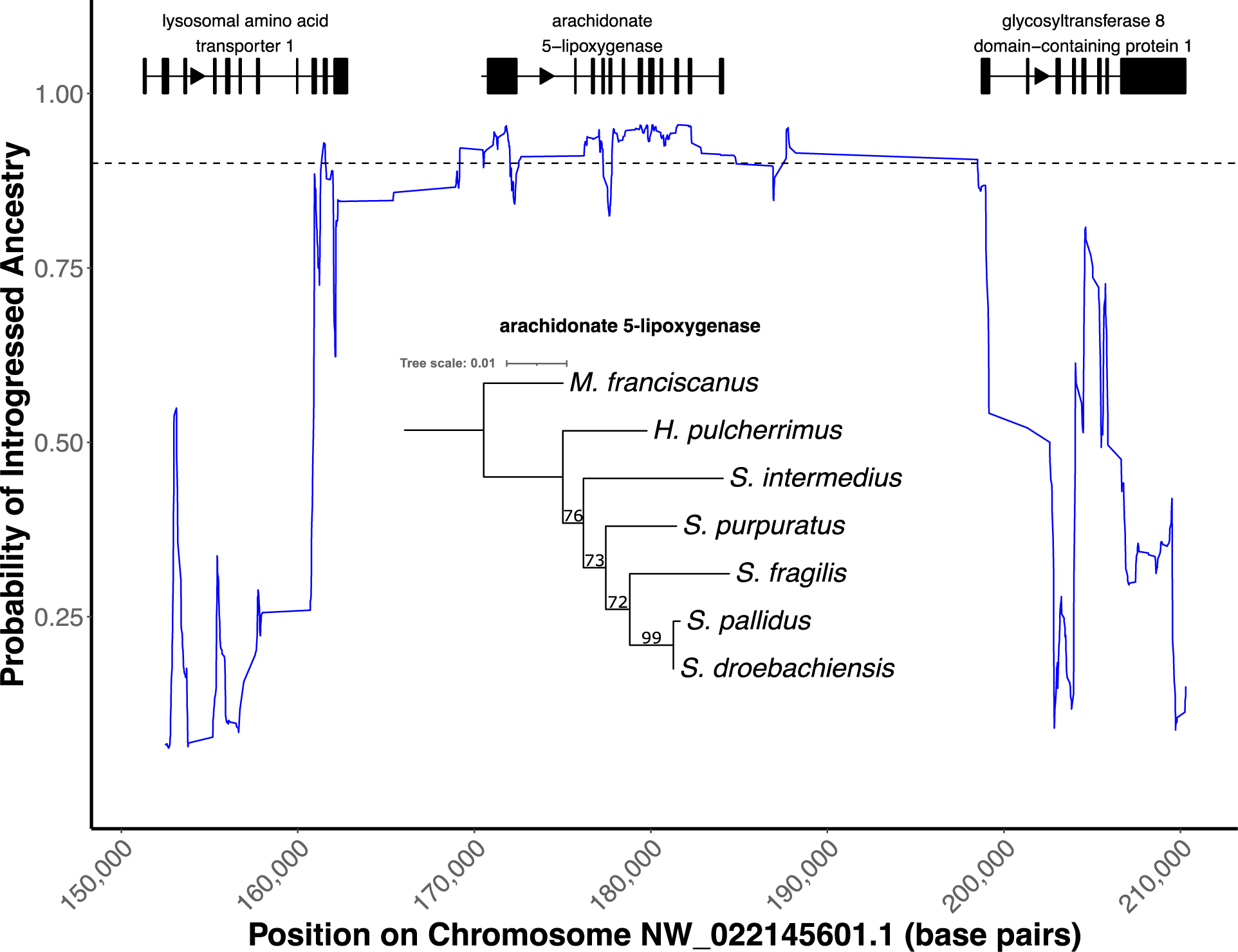
The introgression tract overlapping arachidonate 5-lipoxygenase, a gene with a history of positive selection within the strongylocentrotid sea urchin family. The maximum likelihood gene tree for the longest isoform of arachidonate 5-lipoxygenase shown inside the plot strongly supports the introgression tree topology ((*S. droebachiensis*, *S. pallidus*), *S. fragilis*).

**Table 4.** Summary of the top genes with introgressed bases that had significant branch-sites tests on either the *S. pallidus* or *S. droebachiensis* terminal branches.

| Gene Info |  | Selective History |  | Percent Coding Bases Introgressed |  | Spal - Sdro boot<sup>c</sup> | Mean Coverage Depth |  |  |  |
| --- | --- | --- | --- | --- | --- | --- | --- | --- | --- | --- |
| NCBI LOC ID | Name | PSG<sup>a</sup> | Branch<sup>b</sup> | prob > 0.9 | prob > 0.8 |  | Sdro | Sfra | Spal | Hpul |
| LOC100893326 | kremen protein 1 | yes | *Spal* | 34.9 | 54.7 | 82 | 26 | 35.9 | 9.7 | 19.9 |
| LOC575079 | arylsulfatase | yes | *Spal* | 18.5 | 25.7 | 67 | 34.3 | 47.9 | 17.3 | 38.2 |
| LOC580597 | sodium- and chloride-dependent neutral and basic amino acid transporter B(0+) | yes | *Spal* | 14.5 | 21.1 | 72 | 28 | 28.9 | 10.8 | 31.9 |
| LOC590964 | fibrosurfin-like | yes | *Spal* | 7.1 | 18.3 | 100 | 44.5 | 51.3 | 16.3 | 110.7 |
| LOC105445324 | muscarinic acetylcholine receptor M5-like | no | *Spal* | 0.0 | 39.1 | 87 | 33.1 | 33.8 | 6.2 | 53.8 |
| eef1g | eukaryotic translation elongation factor 1 gamma | yes | *Spal* | 0.0 | 28.6 | 89 | 34.3 | 35.4 | 16.8 | 40.2 |
| LOC577068 | prominin 1 | yes | *Spal* | 0.0 | 12.3 | 96 | 30.7 | 41.3 | 16.1 | 40.2 |
| LOC584837 | PHD finger protein 8 | no | *Sdro* | 20.3 | 42.8 | 100 | 23.2 | 38.3 | 9.6 | 41.7 |
| LOC580943 | structural maintenance of chromosomes 1A | yes | *Sdro* | 4.3 | 39.7 | 59 | 25 | 35.7 | 11.7 | 37.5 |
| LOC105442321 | toll-like receptor 3 | no | *Sdro* | 0.0 | 13.3 | 94 | 58.1 | 54.8 | 18.6 | 73.7 |
| LOC575208 | death-inducer obliterator 1 | no | *Sdro* | 0.0 | 10.0 | 90 | 41.2 | 41.8 | 11.6 | 46.4 |

<sup>a</sup>PSG - Positively Selected Gene. A gene with a significant sites tests of positive selection across the strongylocentrotid family.
<sup>b</sup>Branch with significant branch-sites test.
<sup>c</sup>Bootstrap support for the branch grouping *S. pallidus* and *S. droebachiensis* as sister taxa
Species abbreviations: *Sdro*, *S. droebachiensis*; *Sfra*, *S. fragilis*; *Spal*, *S. pallidus*; *Hpul*, *H. pulcherrimus*.

To determine whether any additional classes of genes were over- or underrepresented in the introgressed set, we tested the genes with more than half of their bases introgressed at the 80% posterior probability threshold for gene ontology enrichment using PANTHER18.0 (Mi et al. 2019; Thomas et al. 2022). Only two significant terms remained after applying a false discovery rate (FDR) correction of 5%. There was an under-enrichment of the cellular component terms “plasma membrane” (GO:0005886, p = 0.014) and “cell periphery” (GO:0071944, p = 0.002). Interestingly, the cellular component term “membrane” (GO:0016020) was enriched in the set of genes with histories of positive selection from Kober and Pogson (2017). Furthermore, the molecular function term “calcium ion binding” (GO:0005509) and biological process term “proteolysis” (GO:0006508) were overrepresented in the set of positively selected genes (Kober and Pogson 2017) and underrepresented in the collection of introgressed genes, though not significant after correction.

## Discussion

Here, we characterized the genomic landscape of introgression between two sea urchin species to gain insight into the factors determining the fate of introgressed variation and the behavior of selection following introgression. Our study is among the first to perform local ancestry inference with whole-genome sequencing data in a high gene flow marine invertebrate group. The strongylocentrotid sea urchin family stands out relative to other population genetic models for their massive effective population sizes, highly efficient selection, and high gene flow across ocean basins. Although the species are well-diverged (4.2 - 19.0 mya), and natural hybrids are rare, many of the species show strong signals of historical introgression (Glasenapp and Pogson 2023). In our analysis of introgression between *S. pallidus* and *S. droebachiensis*, we found strong evidence for genome-wide selection against introgression, including two chromosomes depleted of introgression warranting further examination. Although our results suggest that slowly evolving loci with low divergence are more likely to be able to move between species, introgression has also likely been an important source of adaptive genetic variation. Between 1% and 6% of coding bases supported introgression, and numerous genes with histories of positive selection also had a significant number of introgressed coding bases. A handful of the introgressed genes with histories of selection are involved in defense, including arachidonate 5-lipoxygenase, glutathione peroxidase, and toll-like receptor 3. Additionally, the introgression of many hyalin genes distributed across multiple chromosomes suggests potential functional and adaptive significance, possibly related to defense. Hyalin is a large glycoprotein and a major component of the hyaline layer, the egg extracellular matrix that serves as a cell adhesion substrate during gastrulation (McClay and Fink 1982; Adelson and Humphreys 1988; Wessel et al. 1998). Kober and Pogson (2017) have suggested that the prevalence of positive selection at membrane or extracellular proteins (such as collagens) might be driven by pathogen defense.

Consistent with theoretical predictions about the retention of introgressed ancestry, we found introgression to be more common in genomic regions expected to be under weaker selection. These regions exhibited lower gene density, reduced divergence, slower rates of evolution, and fewer positively selected genes than the non-introgressed genome-wide background. We find it unlikely that these patterns were driven by increased power to detect introgression in low divergence regions, as PhyloNet–HMM requires sequence divergence to detect introgression (Schumer et al. 2016). Furthermore, a negative correlation between introgressed ancestry and sequence divergence has been observed in humans (Vernot and Akey 2014), and many studies have found depleted introgression in functional regions (Teeter et al. 2008; Sankararaman et al. 2014; Janoušek et al. 2015; Juric et al. 2016; Sankararaman et al. 2016; Schumer et al. 2016; Maxwell et al. 2018; Petr et al. 2019). Reduced introgression in regions with high divergence or functional density is likely explained by divergent regions harboring loci underlying local adaptation or Dobzhansky-Muller incompatibilities (Moran et al. 2021). Unfortunately, limited information about natural hybrids and ecological selection in the strongylocentrotid family precludes distinguishing between the different sources of selection against introgressed variation.

Our findings are at odds with those of Schumer et al. (2016), who characterized introgressed *Xiphophorus cortezi* ancestry in *X. nezahualcoyotl* genomes and found that introgressed regions had higher sequence divergence, gene density, and rates of synonymous and nonsynonymous substitutions than the genome-wide background. They demonstrated that the higher divergence of introgressed regions was likely driven by introgression at genes not under strong selective constraint, which is still consistent with genome-wide selection against introgression. The unique results in the *Xiphophorus* system may be driven by the fact that recombination hotspots are concentrated near promoter-like features in swordtails and occur further from transcription start sites in humans and other species with PRDM9-direction recombination (Myers et al. 2005; Coop et al. 2008; Baker et al. 2017; Moran et al. 2021). Furthermore, the *X. nezahualcoyotl* swordtail genome samples had low genetic diversity (*θπ*: 0.00025 - 0.00082), indicating that low or fluctuating effective population sizes and less efficient selection could have also contributed to the higher-than-expected amount of introgression in gene dense regions. The differences between our findings and those of Schumer et al. (2016) highlight the importance of characterizing admixture and introgression across more taxonomic groups.

We believe our estimates of the proportion of the genome introgressed were conservative for several reasons. First, introgression between *S. pallidus* and *S. droebachiensis* is likely historical, making it harder to detect because recombination breaks introgressed haplotypes into progressively shorter tracts over time, and new mutations obscure the history of introgression. Most of the strongylocentrotid genes that showed introgression were not fully introgressed, especially those with histories of positive selection. Instead, many genes had one or more small regions with very strong support for introgression. Due to the old age of introgression and the high expected amount of recombination in strongylocentrotid urchins, the scale of introgression is likely at the exon level rather than the whole gene level. Detecting introgressed regions at this small scale is a major challenge for any statistical method, given the limited number of variants in an individual exon. Second, randomly resolving heterozygous sites to create multiple sequence alignments causes switching between maternal and paternal chromosomes, fragmenting introgressed haplotypes that are heterozygous in our samples and biasing our detection towards introgressed variation that has been fixed. When an introgression tract is heterozygous for ancestry, switching between introgressed and non-introgressed ancestry may lead to ambiguous posterior probabilities (Schumer et al. 2016). For this reason, the introgression tracts we detected are likely fixed and old.

Theory predicts a positive correlation between the extent of introgression and the local recombination rate (Veller et al. 2023). Unfortunately, information on recombination in the strongylocentrotid sea urchins is extremely limited, preventing us from testing this relationship. An outstanding question remains whether differences in recombination rates drove the differences in introgression among the different strongylocentrotid chromosomes. If the number of crossovers per meiosis is constant among chromosomes, shorter chromosomes should have higher per-base recombination rates and retain more introgressed variation. However, we did not find a significant relationship between introgression and chromosome length, and the smallest chromosome had the least amount of introgression, which is inconsistent with expectations.

Without polymorphism data for *S. pallidus* and *S. droebachiensis*, we can only speculate about the proportion of introgression tracts driven to fixation by positive selection. However, selection is expected to be very efficient in these sea urchin species. For example, it was conservatively estimated that 15% of strongylocentrotid single-copy orthologs had experienced positive selection (Kober and Pogson 2017) and *S. purpuratus* shows selection on preferred usage of synonymous codons (Kober and Pogson 2013). Furthermore, the introgression tracts documented in this study are likely historical and fixed. *S. pallidus* and *S. droebachiensis* diverged 5.3 - 7.6 million years ago (Kober and Bernardi 2013) and natural hybrids between the two are rarely observed (Vasseur 1952). Given the likely old age of introgression, the high expected efficiency of selection, and our bias toward detecting high-frequency variants, it is not unreasonable to assume that a small proportion of the introgression tracts spanning coding regions contained advantageous mutations. Future studies will test for recent selection at the genomic intervals inferred to have been introgressed and look for adaptive introgression in promoter regions upstream of genes.

In summary, our study documented strong evidence for genome-wide selection against introgressed variation, suggesting that slowly evolving, low-divergence genomic regions are more likely to move between species and avoid negative selection following hybridization and introgression. However, despite strong selection against introgression, we also identified numerous candidate adaptively introgressed genes, suggesting that introgression has been an important source of adaptive genetic variation. The strongylocentrotid sea urchin family represents a valuable model system for further characterization of introgression given the high amount of gene flow and genetic diversity among the different species.

## Materials and Methods

### Study System

Four species forming a rooted triplet were used in the present study: (*Hemicentrotus pulcherrimus*, (*Strongylocentrotus pallidus*, (*S. droebachiensis*, and *S. fragilis*))). The metadata for the sample accessions are presented in Supplementary Table S1. The three *Strongylocentrotus* taxa were sampled from the East Pacific: *S. droebachiensis* and *S. pallidus* were dredged from Friday Harbor, WA, and *S. fragilis* was collected in Monterey Bay. *H. pulcherrimus* was chosen as the outgroup as it was sampled from the West Pacific (coastal Japan by Y. Agatsuma) and diverged from the *Strongylocentrotus* taxa 10 – 14 mya (Kober and Bernardi 2013). *S. pallidus* and *S. droebachiensis* have broad, overlapping Holarctic distributions with ample opportunity for hybridization. They co-occur in the West Pacific, East Pacific, Arctic, West Atlantic, and East Atlantic Oceans. The geographic history of speciation is challenging to interpret, but fossil evidence confirms that both species speciated in the Pacific and crossed the Bering Sea in the late Miocene to colonize the Arctic and Atlantic Oceans (Durham and MacNeil 1967). Both species show little differentiation between the Pacific and Atlantic due to high trans-Arctic gene flow (Palumbi and Kessing 1991; Addison and Hart 2005; Addison and Kim 2022).

The eggs of *S. droebachiensis* are highly susceptible to fertilization by heterospecific sperm (Strathmann 1981; Levitan 2002), and hybrid matings readily occur when spawning *S. droebachiensis* females are closer to heterospecific males than conspecific males (Levitan 2002). Hybrids between *S. pallidus* and *S. droebachiensis* have also been successfully reared and backcrossed in the lab (Strathmann 1981). Although reproductive isolation between *S. pallidus* and *S. droebachiensis* appears incomplete, the two species remain distinct across their overlapping ranges (Vasseur 1952; Strathmann 1981). However, hybrids of *S. pallidus* and *S. droebachiensis* morphologically resemble *S. pallidus* as larvae and *S. droebachiensis* as adults, so the frequency of natural hybrids may be underestimated. Introgression between *S. pallidus* and *S. droebachiensis* has been previously detected (Addison and Hart 2005; Harper et al. 2007; Addison and Pogson 2009; Pujolar and Pogson 2011; Glasenapp and Pogson 2023).

### Data Pre-Processing

A single genome from each strongylocentrotid species had been previously sequenced on the Illumina HiSeq 2500 (Kober and Bernardi 2013; Kober and Pogson 2017). The raw sequencing reads were pre-processed following GATK’s Best Practices (Van der Auwera et al. 2013). Briefly, adapters were marked with Picard MarkIlluminaAdapters, and sequencing reads were mapped to the *S. purpuratus* reference genome (Spur_5.0) with bwa-mem2 v2.2.1 (Vasimuddin et al. 2019). Duplicates were marked with Picard MarkDuplicates, and reference mapping was evaluated using samtools flagstat (Danecek et al. 2021) and mosdepth v0.3.3 (Pedersen and Quinlan 2018). Variant calling and joint genotyping were performed with GAKT’s HaplotypeCaller and GenotypeGVCFs. Variants were hard-filtered for skewed values across all samples following GATK recommendations (Caetano-Anolles 2023). Further filtering was done for genotypes with low-quality scores (GQ<20) and low read depth (DP<3), and single nucleotide variants (SNVs) within three base pairs of an indel were excluded.

### PhyloNet-HMM

We used an updated version of PhyloNet-HMM called PHiMM (Wuyun et al. 2019) to identify genomic regions supporting introgression between *S. pallidus* and *S. droebachiensis*. PhyloNet-HMM is a hidden Markov model that detects breakpoints between regions supporting different phylogenetic relationships, accounting for incomplete lineage sorting (ILS) by allowing for changes in gene trees within both the species and introgression trees (Liu et al. 2014; Schumer et al. 2016). PhyloNet-HMM walks across each chromosome, locates changes in the underlying genealogy, and outputs posterior probabilities for each SNV site, reflecting the likelihood that the site evolved along the species and introgression trees. PhyloNet-HMM has been used to detect introgressed regions in swordtails (Schumer et al. 2016; Powell et al. 2020), house mouse (Liu et al. 2015), North American admiral butterfly (Mullen et al. 2020), *Danaus* butterfly (Aardema and Andolfatto 2016), and snowshoe hare (Jones et al. 2020) genomes. Schumer et al. (2016) conducted performance tests of PhyloNet-HMM on simulated swordtail data and concluded that the approach could accurately distinguish between ILS sorting and hybridization.

Multiple sequence alignments of single nucleotide variant (SNV) sites were created for the 21 largest *S. purpuratus* scaffolds using vcf2phylip (Ortiz 2019). The 21 largest scaffolds correspond to the 21 *S. purpuratus* chromosomes (2n=42) and represent 90% of the 921,855,793 bases in the *S. purpuratus* reference genome assembly (Spur_5.0). Although it is known that *S. purpuratus* has a genetically based sex determination, sex chromosomes have yet to be identified (Pieplow et al. 2023). PhyloNet-HMM only allows for DNA base characters in the input sequence alignments, so all indels were excluded, and only variable sites where all four samples had genotypes passing filter were used. Heterozygous genotypes were randomly resolved because PhyloNet-HMM does not support IUPAC ambiguity codes, and accurate phasing could not be performed across entire chromosomes with a single diploid genome per species and no reference panel (Bukowicki et al. 2016). We ran PhyloNet-HMM on each chromosome 100 times using the default settings to avoid the effects of reaching local optima during hill climbing and averaged the posterior probabilities across independent runs. The average distance between variable sites in the alignments was 85 base pairs. The multiple sequence alignments including invariant sites had 2,361 gaps greater than 25 kb in length, with the largest gap being 585,224 base pairs.

We used two different posterior probability thresholds to identify introgression tracts: 90% and 80%. Introgression tracts were inferred by recording the genomic coordinates of consecutive sites with posterior probabilities at or above the threshold. The mean coverage depth for each introgression tract for each species was calculated with mosdepth (Pedersen and Quinlan 2018), and introgression tracts where any species had coverage depth less than 5x or greater than 100x were excluded. Introgression tracts overlapping a gap of 25 kb or more between adjacent genotypes (including invariant sites) were identified using bedtools intersect (Quinlan and Hall 2010) and trimmed to remove the gap. Coverage depth metrics were estimated for all introgression tracts passing filter using mosdepth (Pedersen and Quinlan 2018). For each introgression tract, overlapping genes and coding bases were recorded using the gff file from the *S. purpuratus* assembly and bedtools intersect (Quinlan and Hall 2010). We also intersected the introgression tracts with the set of genes with a history of positive selection within the strongylocentrotid family identified by (Kober and Pogson 2017).

### Properties of Introgressed Regions

To characterize the genomic landscape of introgression, we compared estimates of absolute nucleotide divergence (*d*_XY_), gene density, coding base density, and rates of evolution (*d*_N_, *d*_S_, and *d*_N_/*d*_S_) for the set of introgressed intervals greater than 10 kb in length to estimates for the non-introgressed genome-wide background (i.e., species tree regions).

#### Divergence

We compared the mean absolute nucleotide divergence (*d*_XY_) of introgression tracts to the mean *d*_XY_ of a random sample of non-introgressed genomic regions of the same number and size as the introgressed intervals. To avoid the confounding effect of introgression between *S. pallidus* and *S. droebachiensis* on *d*_XY_ estimates, we used *H. pulcherrimus* and *S. fragilis*, who have experienced little-to-no introgression. The two species involved in introgression (*S. pallidus*, *S. droebachiensis*) were not included in the divergence measures because introgression from *S. droebachiensis* into *S. pallidus* would decrease *S. fragilis* - *S. pallidus* divergence, introgression from *S. pallidus* into *S. droebachiensis* would decrease *S. fragilis* - *S. droebachiensis* divergence, and introgression between *S. pallidus* and *S. droebachiensis* in either direction would reduce *S. pallidus* - *S. droebachiensis* divergence (see Forsythe et al. 2020).

To obtain distributions of *d*_XY_, we first generated a new genotype (vcf) file for *S. fragilis* and *H. pulcherrimus,* including invariant sites. Vcf files typically only contain variant sites, and *d*_XY_ estimates can be downwardly biased by assuming missing sites are invariant (Korunes and Samuk 2021). We generated the new genotype file by combining the single sample vcf files for *H. pulcherrimus* and *S. fragilis* and performing joint genotyping using GATK’s GenotypeGVCFs with the –include-non-variant-sites option. Variant and invariant sites were then split into separate files for filtering. Variant sites were hard-filtered for skewed values across all samples following GATK recommendations (Caetano-Anolles 2023). The variant and invariant site vcf files were then merged back together, and genotypes with low-quality scores (GQ<30), low read depth (DP<8), or low reference genotype confidence (RGQ<30) were set to missing.

The random sample of non-introgressed regions was created by randomly permuting intervals of the same number and size of the 10kb introgression tracts into regions confidently called for the species tree using bedtools shuffle (Quinlan and Hall 2010). We used pixy (Korunes and Samuk 2021) to calculate *d*_XY_ for each genomic interval in the sets of introgressed and non-introgressed intervals and implemented a bootstrap comparison of means to test for a significant difference between the mean *d*_XY_ of introgressed and non-introgressed regions. We pooled all *d*_XY_ values for both sets, bootstrap resampled the pool in pairs 100,000 times, and calculated the difference in mean *d*_XY_ between bootstrapped pairs to generate the distribution of differences in means expected if there were no difference between mean introgressed *d*_XY_ and mean non-introgressed *d*_XY_. We then compared the true difference in mean *d*_XY_ between the introgressed and non-introgressed regions to the null distribution to calculate a p-value.

#### Rate of Evolution

We next compared the rate of evolution of introgressed genes to that of non-introgressed genes. For *H. pulcherrimus* and *S. fragilis*, we first identified genes with more than half of their bases declared introgressed at each posterior probability threshold, excluding those with mean coverage depth less than 10x or greater than 100x, with fewer than 75% of their coding bases covered by one read or fewer than 50% by ten reads, or with premature stop codons. We next created sequence alignments of *H. pulcherrimus* and *S. fragilis* and for each introgressed gene passing filter using vcf2fasta, and estimated *d*_N_, *d*_S_, and *d*_N_/*d*_S_ using codeml model M0 of PAML (Yang 2007). We specified the cleandata=1 option to remove sites with ambiguity data. To obtain estimates of *d*_N_, *d*_S_, and *d*_N_/*d*_S_ for non-introgressed genes, we identified all genes with more than half of their bases confidently called for the species tree, filtering by the same metrics as the introgressed genes. We then randomly sampled the identified non-introgressed genes to get a sample the same size as the number of introgressed genes and estimated *d*_N_, *d*_S_, and *d*_N_/*d*_S_ for each gene. We compared the means of each metric between introgressed and non-introgressed genes using the same bootstrap comparison of means procedure used in the *d*XY analysis. For each metric (*d*_N_, *d*_S_, and *d*_N_/*d*_S_), all values from both the introgressed and non-introgressed sets were pooled. The pool was bootstrap resampled in pairs 100,000 times, and the difference in means for each metric was calculated between bootstrapped pairs to generate the distribution of differences in means expected if there were no difference between mean introgressed *d*_N_, *d*_S_, and *d*_N_/*d*_S_ and mean non-introgressed *d*_N_, *d*_S_, and *d*_N_/*d*_S_. We then compared the true difference in means between the introgressed and non-introgressed regions to the null distribution to calculate a p-value.

#### Gene Density

To determine whether introgressed regions were more or less likely to overlap protein-coding genes, genes with a history of positive selection, and coding bases than the genome-wide background, we bootstrap resampled the introgression tracts with replacement to create 1,000 pseudoreplicate datasets. For each, we counted the number of overlapping coding bases, the number of protein-coding genes with more than half of their bases declared introgressed, and the number of genes with a history of positive selection identified by Kober and Pogson (2017). We identified the overlapping genes and coding bases by intersecting the introgression tract interval files with the protein-coding gene and CDS coordinates for *S. purpuratus* using bedtools intersect. To standardize the protein-coding gene counts, we divided the values by the total length of the interval files in megabases. To normalize the coding base counts, we divided the number of coding bases by the total number of bases in the interval file. To generate null distributions representative of the genome-wide background for protein-coding genes, genes with a history of positive selection, and coding base counts, we created 1,000 replicate interval sets by randomly permuting intervals of the same number and size of the 10kb introgression tracts into regions confidently called for the species tree using bedtools shuffle (Quinlan and Hall 2010). We then compared the mean and standard deviation of each metric for the introgressed set to the 95% confidence intervals of the null distribution representative of the genome-wide background.

## Data Accessibility and Benefit-Sharing

### Data Accessibility Statement

The data and code supporting this study’s findings are available on Dryad (https://doi.org/10.5061/dryad.fn2z34v1k). Raw sequence reads are available in the NCBI SRA (BioProject PRJNA391452).

## Acknowledgments

Funding for data collection for the study was provided by the National Science Foundation (DEB-1011061), the STEPS Foundation, Friends of Long Marine Lab, and the Myers Trust. The funding bodies did not participate in research design, sample collection, data analysis, or manuscript writing. We thank UC Santa Cruz for access to the Hummingbird computational cluster and Qiqige Wuyun for their help with PhyloNet-HMM. We thank Matthew Kustra for his help with data visualization and PAML. We thank Kord Kober, Pete Raimondi, and Russ Corbett-Detig for their helpful comments on the manuscript. We thank two anonymous reviewers for their constructive feedback that greatly improved the manuscript. The authors have no conflicts of interest to declare.

