## Supplementary Figure S1 for "Selection shapes the genomic landscape of introgressed ancestry in a pair of sympatric sea urchin species"

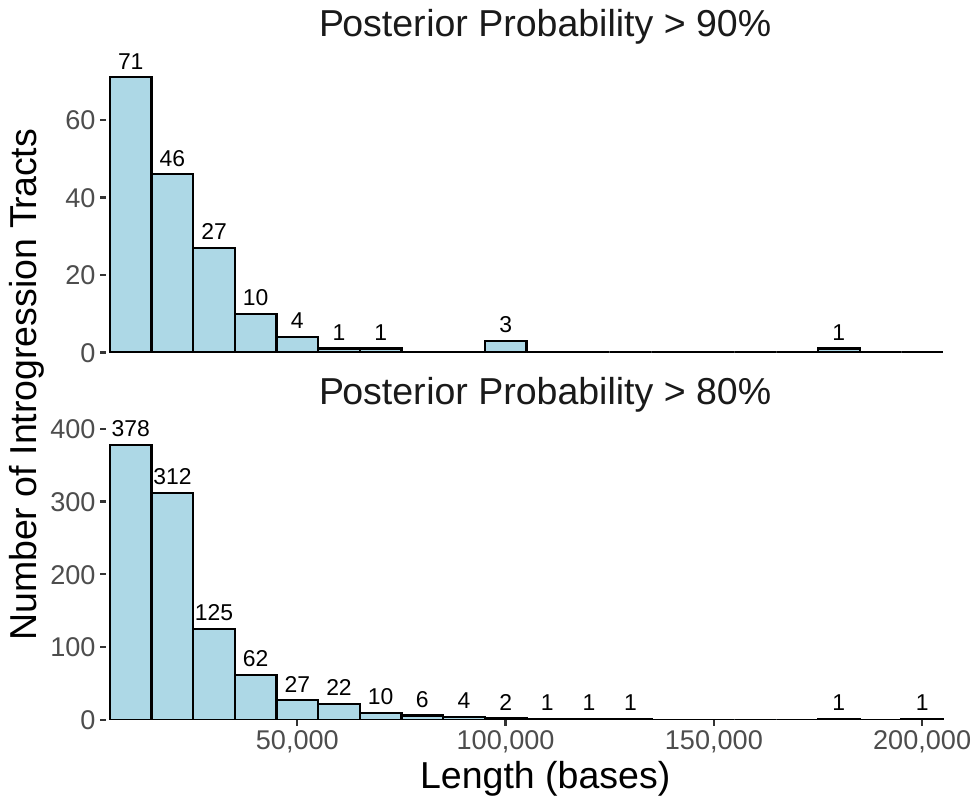

**Supplementary Figure S1.** Distribution of introgression tract lengths for introgression tracts greater than 10 kb. Top panel: tract length distribution using a posterior probability cutoff of 90% as evidence of introgression. Bottom panel: tract length distribution using a posterior probability cutoff of 80%.

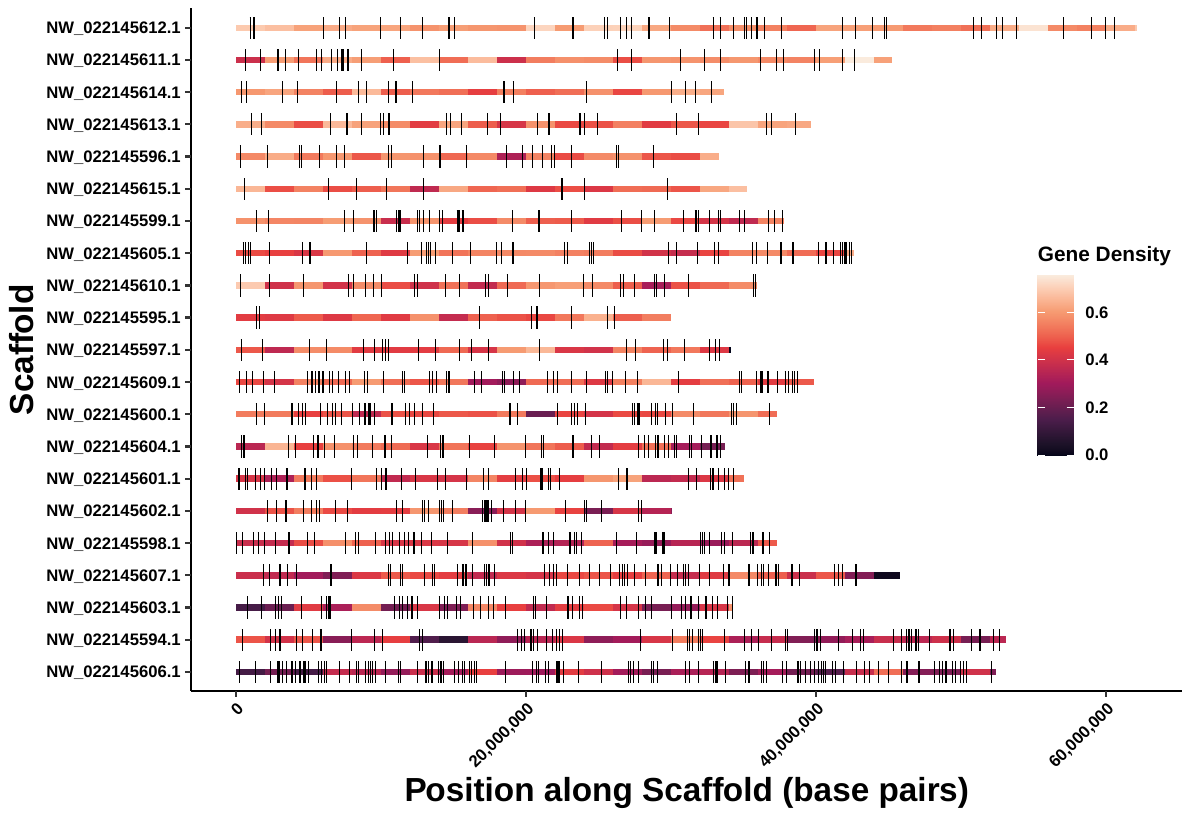

**Supplementary Figure S2**. The 953 introgression tracts greater than 10 kb in length by chromosome (posterior probability > 80%). The introgression tracts are displayed as black rectangles along the chromosomes. The chromosomes are ordered by gene density (descending). The chromosomes are colored by gene density in windows of 2Mb.

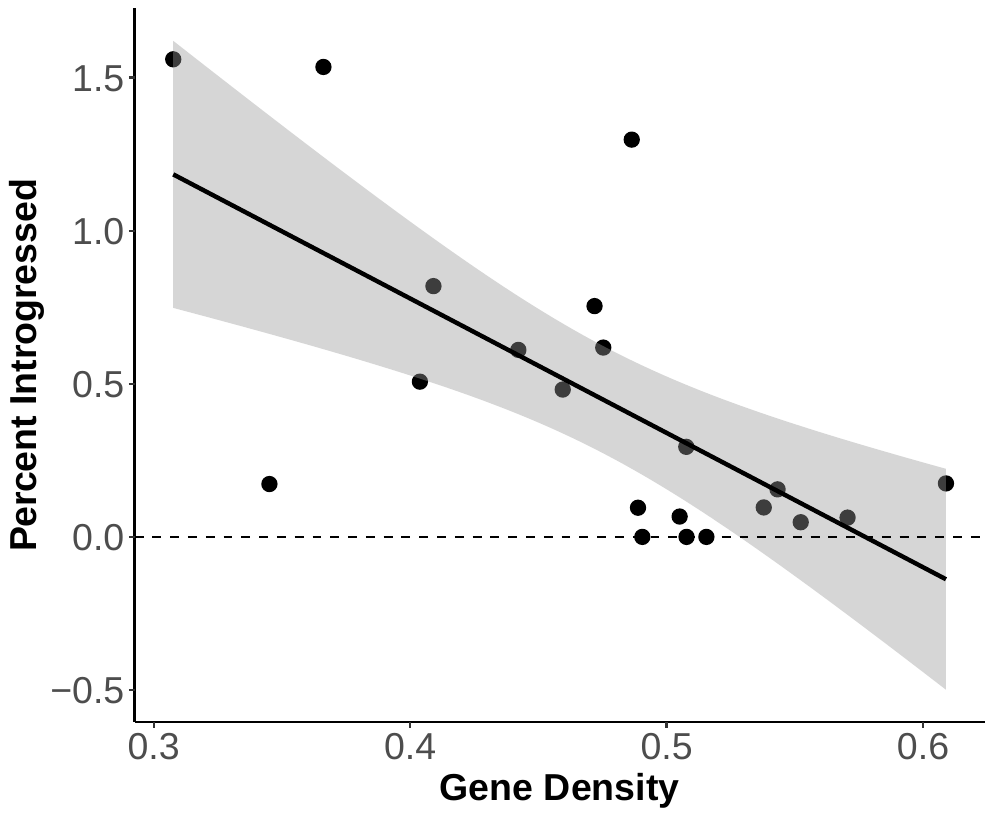

**Supplementary Figure S3.** Percent chromosome introgressed by chromosome gene density. Chromosome gene density was calculated as the proportion of bases falling within genic regions (including introns and untranslated regions) for each chromosome. Percent introgressed was calculated as the total length of the 10 kb introgression tracts for each chromosome divided by the total length spanned in the chromosome multiple sequence alignments scanned by PhyloNet-HMM.

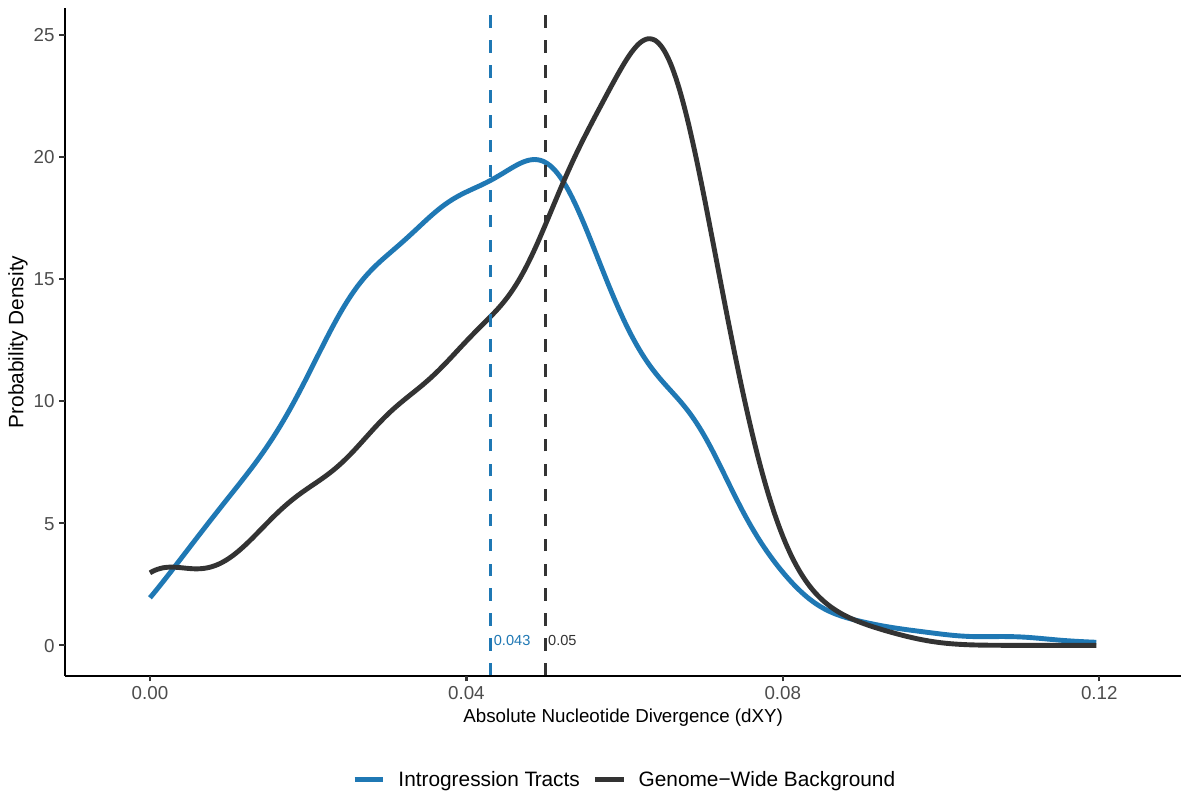

**Supplementary Figure S4**. Absolute nucleotide divergence (*d*_XY_) for the introgressed intervals at the 80% posterior probability threshold vs. a random sample of non-introgressed intervals of the same number and length confidently called for the species tree by PhyloNet-HMM.

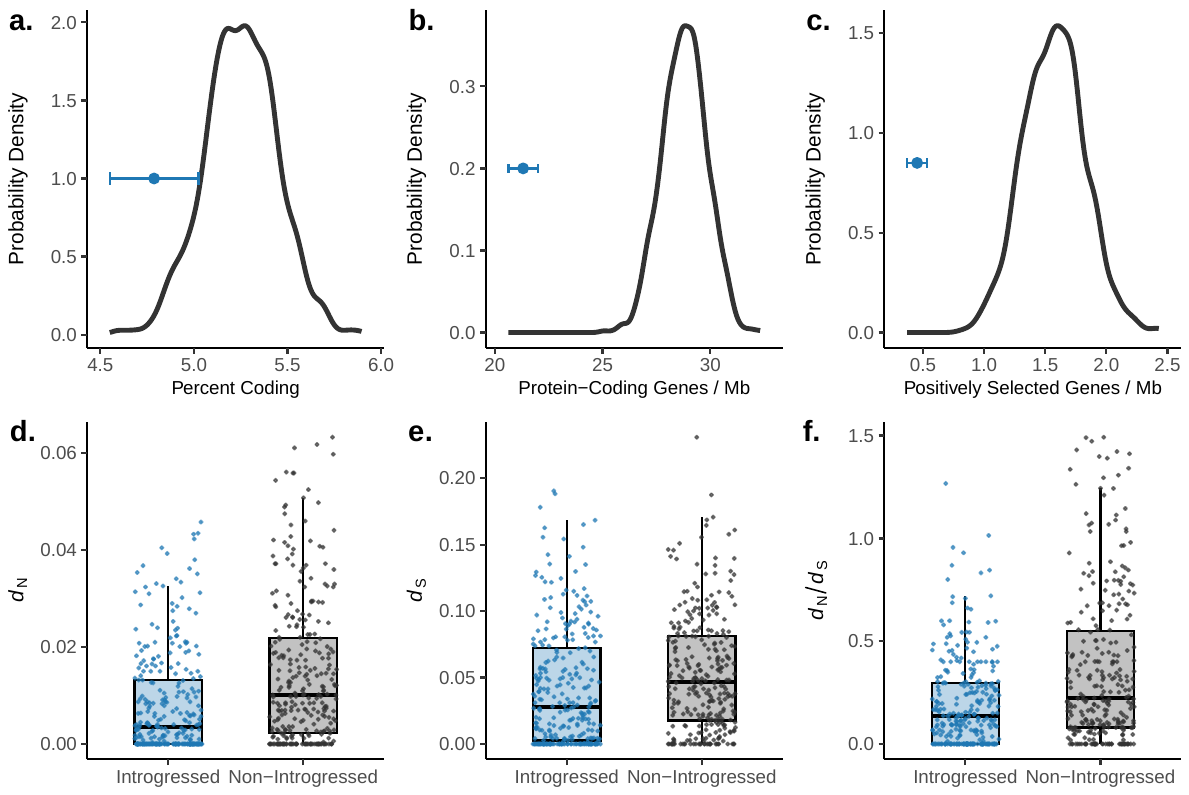

**Supplementary Figure S5.** Figure 4. Properties of introgressed regions and genes at the 80% posterior probability threshold relative to random non-introgressed genes representative of the genome-wide background. (a) The percentage of bases that are coding for the introgression tracts (blue) is lower than the genome-wide background. (b) The number of overlapping protein coding genes, standardized by the combined number of introgressed bases in Mb, is lower than the genome-wide background. (c) The number of overlapping positively selected genes, standardized by the number of bases in the interval files, is lower for introgression tracts than the genome-wide background. Errors bars in (a-c) represent the standard deviation. (d-f) *d*_N_, *d*_S_, and *d*_N_/*d*_S_ are lower for introgressed genes than non-introgressed genes.
